# X-Atlas/Orion: Genome-wide Perturb-seq Datasets via a Scalable Fix-Cryopreserve Platform for Training Dose-Dependent Biological Foundation Models

**DOI:** 10.1101/2025.06.11.659105

**Authors:** Ann C Huang, Tsung-Han S Hsieh, Jiang Zhu, Jackson Michuda, Ashton Teng, Soohong Kim, Elizabeth M Rumsey, Sharon K Lam, Ikenna Anigbogu, Philip Wright, Mohamed Ameen, Kwontae You, Christopher J Graves, Hyunsung John Kim, Adam J Litterman, Rene V Sit, Alex Blocker, Ci Chu

## Abstract

The rapid expansion of massively parallel sequencing technologies has enabled the development of foundation models to uncover novel biological findings. While these have the potential to significantly accelerate scientific discoveries by creating AI-driven virtual cell models, their progress has been greatly limited by the lack of large-scale high-quality perturbation data, which remains constrained due to scalability bottlenecks and assay variability. Here, we introduce “Fix-Cryopreserve-ScRNAseq” (FiCS) Perturb-seq, an industrialized platform for scalable Perturb-seq data generation. We demonstrate that FiCS Perturb-seq exhibits high sensitivity and low batch effects, effectively capturing perturbation-induced transcriptomic changes and recapitulating known biological pathways and protein complexes. In addition, we release X-Atlas: Orion edition (X-Atlas/Orion), the largest publicly available Perturb-seq atlas. This atlas, generated from two genome-wide FiCS Perturb-seq experiments targeting all human protein-coding genes, comprises eight million cells deeply sequenced to over 16,000 unique molecular identifiers (UMIs) per cell. Furthermore, we show that single guide RNA (sgRNA) abundance can serve as a proxy for gene knockdown (KD) efficacy. Leveraging the deep sequencing and substantial cell numbers per perturbation, we also show that stratification by sgRNA expression can reveal dose-dependent genetic effects. Taken together, we demonstrate that FiCS Perturb-seq is an efficient and scalable platform for high-throughput Perturb-seq screens. Through the release of X-Atlas/Orion, we highlight the potential of FiCS Perturb-seq to address current scalability and variability challenges in data generation, advance foundation model development that incorporates gene-dosage effects, and accelerate biological discoveries.

## Introduction

The advent of massively parallel sequencing technologies, particularly single-cell RNA sequencing (scRNA-seq), has fundamentally reshaped biological inquiry, offering unprecedented resolution into the heterogeneity and dynamics of cellular systems ^1,2^. This explosion of high-throughput and high-dimensional data has catalyzed the development of sophisticated computational approaches, including the adaptation of large language model architectures into biological “foundation models” ^3–7^. These models hold tremendous promise, potentially enabling the creation of *in silico* representation of cells – AI-driven “virtual cells” – that could accelerate biology discovery and therapeutic development ^8^. By learning complex patterns from vast datasets, foundation models aim to predict cellular responses and uncover novel biological mechanisms.

Despite their power in capturing correlative structures within observational biological data, a critical gap remains: current foundation models often struggle to robustly infer causal relationships and predict the outcome of specific interventions, underperforming simpler models ^9–11^. This limitation stems from the inherent difficulty of discerning cause and effect from passive observations alone ^12,13^. Truly predictive and mechanistic models require data that directly probes the consequences of defined perturbations. To bridge this gap and unlock the full potential of AI in biology, there is a pressing need for large-scale, high-quality datasets derived from systematic perturbation experiments.

Perturb-seq, a technology combining pooled CRISPR screening with scRNAseq, has emerged as a powerful tool for generating such causal data, directly linking genetic perturbations to their transcriptomic consequences at single-cell resolution ^14–16^. By systematically knocking down or activating genes and observing the resulting changes in gene expression, Perturb-seq provides the rich, interventional data necessary to train models capable of understanding and predicting causal biological mechanisms. While pioneering studies have demonstrated the feasibility of genome-scale Perturb-seq experiments ^17–20^, significant hurdles remain in scaling these efforts further. Current protocols often face substantial logistical challenges, including the need to process fresh cells, limitations in throughput, and batch-to-batch variability. These factors introduce technical noise that can confound biological signals and compromise the quality of data needed for training robust, generalizable foundation models.

To address these critical bottlenecks, we developed “Fix-Cryopreserve-ScRNAseq” (FiCS) Perturb-seq, an industrialized platform for scalable, consistent, and high-quality Perturb-seq data generation. We describe the FiCS Perturb-seq methodology and demonstrate its ability to produce sensitive transcriptomic data with markedly reduced batch effects compared to existing approaches. In addition, we release X-Atlas: Orion edition (X-Atlas/Orion), a significant resource comprising two genome-wide FiCS Perturb-seq datasets in HCT116 and HEK293T cells. This dataset, the largest publicly available Perturb-seq resource to our knowledge, serves both as to validate the FiCS Perturb-seq platform and as a foundational dataset, poised to fuel new analytical approaches in computational biology. Indeed, the scale and depth of X-Atlas/Orion allow for advancements beyond common analytical paradigms. For example, current approaches in Perturb-seq computational analysis, including the training of many biological foundation models, treat genetic perturbations as categorical variables (e.g., ‘on’ vs. ‘off’), potentially overlooking the continuous spectrum of perturbation strengths and the corresponding dose-dependent cellular responses richly present in the data. Leveraging the volume and quality of data within X-Atlas/Orion, we enable the study of dose-dependent genetic effects using sgRNA abundance as a quantitative proxy for perturbation strength. By treating perturbation as a continuous variable, our approach offers a more refined framework to enhance the predictive power and biological insight of future causal models.

## Results

### Establishing the FiCS Perturb-seq Platform

A Perturb-seq experiment typically involves engineering cell lines for stable Cas9 expression, delivering CRISPR sgRNA libraries via lentivirus, preparing and sequencing single-cell libraries, and performing computational analysis (**Figure 1A**). However, scaling Perturb-seq to genome-wide experiments presents several challenges: 1) Most single-cell platforms are optimized for freshly harvested cells, where handling induced cellular stress can confound transcriptomics phenotyping – an issue exacerbated by the throughput mismatch between genome-wide perturbations (tens of millions of input cells) and platforms like 10x Genomics GEM-X (∼200,000 cells per batch). 2) In small-scale experiments, fluorescence-activated cell sorting (FACS) enrichment for viable and perturbed cells enhances efficiency, but for genome-wide studies, FACS processing is prohibitively slow and can induce substantial cellular stress or death. 3) The reliance on fresh cells prevents a separate upfront quality control (QC) on a subset of samples, requiring full commitment to an expensive, labor-intensive experiment at the time of cell harvest. 4) Downstream library preparation involves hundreds of individual reactions, where automation is crucial for improving efficiency and minimizing human error and variability.

**Figure 1.**
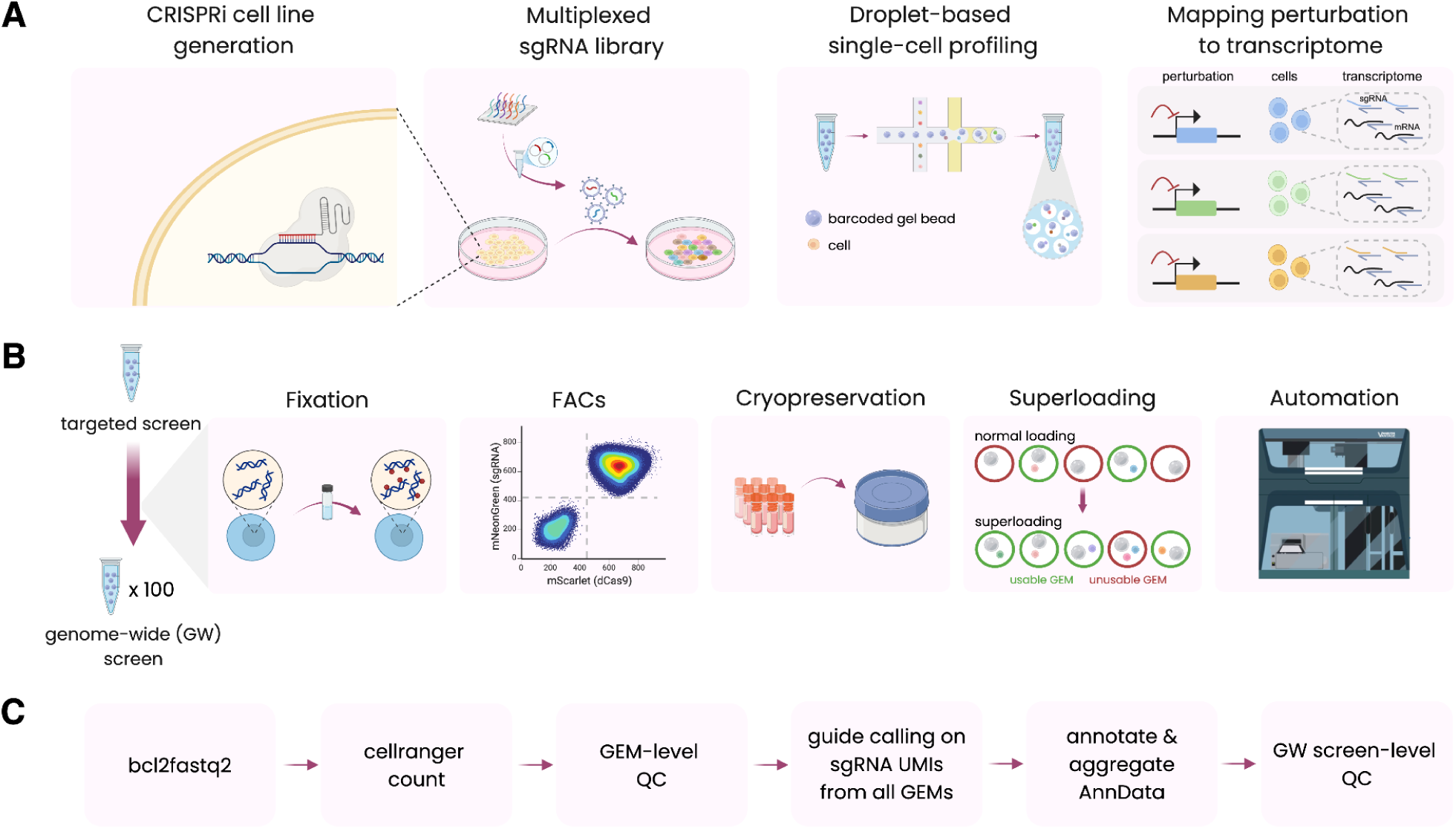
Industrialized Perturb-seq Platform Workflow. **A**. Overview of a CRISPRi Perturb-seq screen, which involves generating stable CRISPRi cell lines, delivering pooled sgRNA libraries targeting genes of interest, preparing single-cell libraries, and mapping perturbations to transcriptomic profiles through computational analysis. **B**. Overview of the FiCS Perturb-seq platform. Cells are DSP-fixed, sorted to isolate viable, dCas9 and sgRNA-positive populations, cryopreserved, superloaded during single-cell library preparation, and sequenced from libraries prepared using automation, which enables scale-up to genome-wide screens. **C**. Computational pipeline for processing raw sequencing data to GEM-level count matrices, annotating cells with their sgRNA identities, and aggregating GEM-level outputs into a single AnnData object. This AnnData object contains the perturbation of a cell and its corresponding counts matrix for all cells in the screen. Created in BioRender.

To overcome these challenges, we developed an industrialized FiCS Perturb-seq platform with five key workflow optimizations: 1) We chemically preserve cells immediately after dissociation using Lomant’s Reagent (dithiobis(succinimidyl propionate), or DSP), a reversible crosslinker compatible with scRNAseq workflows ^21–24^, minimizing biological and technical noise from subsequent handling while preserving the correspondence of resulting transcriptomes with those from fresh cells (**Figure S1A**). 2) Fixation can alter cell morphology, making live-cell sorting via forward and side scatter unreliable; therefore, we introduce a fixative compatible viability stain (Zombie dye) to aid live-cell selection (**Figure S1B**) while simultaneously sorting for Cas9 and sgRNA dual-expressing cells, enriching for high-quality perturbed cells prior to single-cell analysis (**Figure S1C**). 3) We enable long-term cryopreservation of fixed cells in Bambanker medium supplemented with RNase inhibitor (**Methods**). 4) We “superload” microfluidics channels up to 100k cells / channel (GEM-X 5’ kits, 10x Genomics) to reduce library prep cost and increase throughput by several fold. 5) We automate library preps on a Hamilton Vantage (**Figure 1B**, **Methods**) to remove operator variability across batches.

To facilitate data analysis, we built a pipeline that performs end-to-end processing from FASTQ to analysis-ready outputs, including basecalling, cell barcode alignment and unique molecular identifier (UMI) quantification, sgRNA-calling on the full dataset, and annotation and aggregation of the final AnnData object. Downstream steps, including knockdown (KD) efficiency calculation and model training, were performed in a sharded manner using an on-disk memory-mapped representation of the same AnnData counts to efficiently handle the large size of the dataset (**Figure 1C**, **Methods**).

### FiCS Perturb-seq generates high-quality genome-wide data

We engineered HCT116 (human colorectal carcinoma) and HEK293T (human embryonic kidney), two cell lines widely used in therapeutic research and drug screening ^25,26^, with dCas9-KRAB-Zim3, which has been reported to be more universally applicable and have higher KD efficiencies ^17,27^. We deliver dual-sgRNA targeting the same gene via a tRNA-based lentiviral delivery system (41,780 sgRNAs targeting 18,903 genes and 1,026 non-targeting control pairs), which we found to reduce recombination rates compared with dual-promoter systems, consistent with reported advantage of the tRNA-based delivery system ^28,29^ (**Figure S1D**). sgRNA are decoded using “direct capture”, where a capture sequence primes reverse transcription of sgRNA instead of PolyA-based priming used in CROP-seq ^16,30^.

We demonstrated the utility of FiCS Perturb-seq by performing genome-wide Perturb-seq in both HCT116 and HEK293T, targeting all human protein-coding genes (n = 18,903 genes). We fixed cells immediately upon dissociation, enriched for viable dCas9 and sgRNA dual-expressing cells by FACS, and cryostored cells at −80°C at 2-2.5 million cells / vial. We thawed 4-8 million cells per batch for scRNAseq library generation on the 10x Genomics GEM-X 5’ Platform. All libraries were sequenced on Illumina NovaSeq X Plus instruments. We demonstrated that DSP fixation did not interfere with scRNAseq and yielded a highly sensitive transcriptome capture (5,387 and 5,871 median genes per cell on HCT116 and HEK293T respectively, **Table 1**). Both genes and transcripts detected per cell compared favorably with the benchmarking dataset from Replogle et al. 2022 ^17^ (**Figure 2A**), indicating good RNA quality and high assay sensitivity. Notably, cryostorage in the presence of RNase inhibitors maintained stable RNA-seq quality for up to 140 days in storage (**Figure 2B**), completely decoupling cell dissociation from single cell library preparation, removing a major throughput bottleneck, and simplifying experimental planning and execution. FiCS Perturb-seq also made it possible to perform a pilot QC scRNAseq on a small subset of cells (321,706 out of 9,886,825 cells, or 3.25%, of the HCT116 screen; 1,084,054 out of 21,550,681 cells, or 5.03%, of the HEK293T screen) for QC, before launching expensive and labor-intensive full genome-wide scRNAseq library preparation (**Figure 2C**). Finally, key library preparation steps were automated on a Hamilton Vantage (**Methods**). scRNA results from automated workflow were highly comparable to manual workflow (Pearson correlation = 0.996, **Figure 2D**) such that it could be used to supplement or replace manual workflow where needed. Taken in aggregate, our industrialized workflow generated a sensitive and homogenous transcriptomics dataset as measured by median UMIs per batch when compared with prior genetic and chemical perturbation atlases, with median UMI counts of HCT116 being 1.68 times as deep as the Replogle K562 Perturb-seq dataset ^17,31^ (median UMIs per cell = 11499), and 8.45 times as deep as the Tahoe-100M chemical perturbation dataset ^17,31^ (median UMIs per cell = 2298) (**Table 1**, **Figure 2E**). Moreover, FiCS Perturb-seq yielded more consistent data with significantly greater batch-to-batch correlation than that of Replogle K562 (Wilcoxon rank-sum test, 5.77e-16 and 2.80e-14 for HCT116 and HEK293T vs. Replogle K562 respectively, **Methods**, **Figure 2F**). As a negative control, cross cell line correlation was calculated using randomly paired batches from HCT116 and Replogle K562, which resulted in lower correlation, as expected (median Spearman correlation: 0.74, **Figure S1E**). Furthermore, the median absolute deviation (MAD) of HCT116 and HEK293T was less than that of Replogle K562 (MAD: 2.87e-3, 3.58e-3, and 9.09e-3 in HCT116, HEK293T, and Replogle K562 respectively), highlighting the consistency and reproducibility of data generated using FiCS Perturb-seq^17^.

**Figure 2.**
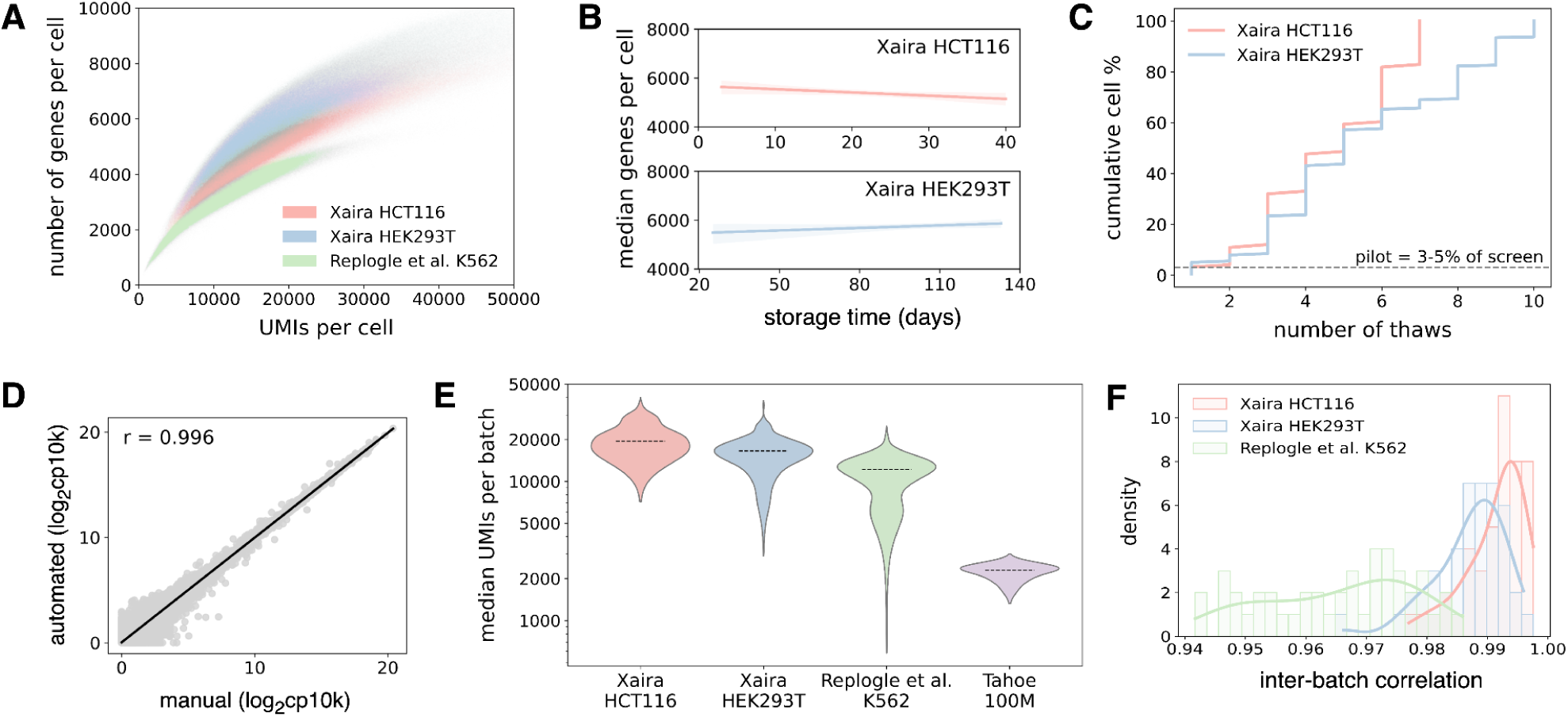
FiCS Perturb-seq delivers sensitive and consistent data. **A**. Scatterplot of UMIs per cell versus number of genes with non-zero counts per cell in the HCT116, HEK293T, and Replogle K562 datasets. **B**. Median genes detected per cell in cryopreserved cells stored up to 40 or 140 days for HCT116 and HEK293T, respectively. **C**. Cumulative cell percentage of screen per cell thaw in HCT116 and HEK293T. 3.25% and 5.03% of cells are used for the initial QC screen in HCT116 and HEK293T, respectively, while the remaining cells are cryopreserved until library preparation. **D**. Correlation between pseudobulk counts from the same cells prepared manually or using the Hamilton Vantage system. Pseudobulk counts are calculated as log2(cp10k + 1), where cp10k represents counts per 10,000, and a pseudocount of 1 is added. **E**. Distribution of median UMIs per cell across GEM groups in HCT116 and HEK293T screens and published genetic and chemical perturbation atlases. Dotted line indicates median UMIs per batch. y-axis is displayed on log10-scale with labels annotated on linear-scale. **F**. Distribution of Spearman correlation coefficients for pairwise pseudobulked gene expression counts from cells containing non-targeting sgRNAs in different batches of the HCT116, HEK293T, and Replogle K562 datasets (median Spearman correlation: 0.993, 0.988, 0.967, respectively).

**Table 1.** QC metrics of the X-Atlas/Orion dataset. # aligned cells, Mean reads per cell, Median UMIs per cell, and Median genes per cell are related to data sensitivity. Median CRISPR UMIs per cell, # cells with two sgRNAs (expected guide pair targeting same gene), Median number of two sgRNA cells per perturbation, and # cells with one sgRNA are related to efficiency of sgRNA capture. Median on-target KD % for two sgRNA cells and Median on-target KD % for one sgRNA cells are primary readouts of CRISPRi efficiency. Percentage of expressed gene perturbations with discernible transcriptomic phenotypes (q-val < 0.05) is related to quantification of secondary effects from perturbation. For details on each metric, see **Methods** section.

|  | HCT116 | HEK293T |
| --- | --- | --- |
| # GEM groups | 109 | 223 |
| # aligned cells | 9,886,825 | 21,550,681 |
| Mean reads per cell | 54,826 | 63,426 |
| Median UMIs per cell | 19,416 | 16,557 |
| Median genes per cell | 5,387 | 5,871 |
| Median CRISPR UMIs per cell | 950 | 184 |
| # cells with two sgRNAs (expected pair) | 3,409,169 | 4,534,299 |
| Median number of two sgRNA cells per perturbation | 141 | 188 |
| Median on-target KD % for two sgRNA cells | 75.4% | 51.5% |
| # cells with one sgRNA | 1,290,167 | 2,326,883 |
| Median on-target KD % for one sgRNA cells | 73.3% | 45.3% |
| Percentage of expressed gene perturbations with discernible transcriptomic phenotypes (q-val < 0.05) | 48.36% | 40.37% |

Not every cellular barcode contained the desired single-target perturbation. This was due to several factors, including the Poisson distribution of lentiviral delivery, high multiplet rates from superloading 10x Genomics channels, and inefficiency in sgRNA calling algorithms. From approximately 30 million sequenced barcodes, we identified around eight million cells harboring two sgRNAs targeting the same intended gene, making the “Orion” dataset the biggest publicly deposited Perturb-seq atlas. This yield of cells with the desired dual sgRNA configuration is consistent with superloading outcomes reported in Nourreddine et al. 2024 ^19^. An additional 3.6 million cells contained only one detectable sgRNA, likely reflecting sensitivity limits in sgRNA-calling algorithms. Notably, these single-sgRNA cells displayed similar KD efficiencies compared with their two-sgRNA counterparts (**Table 1** for full metrics). Despite the potential utility of single-sgRNA cells, we focused subsequent analysis exclusively on the eight million cells carrying the intended sgRNA pairs to ensure stringent filtering.

### FiCS Perturb-seq detects perturbation effects and recovers known biology

We assessed the proportion of perturbations that produced discernible transcriptomic effects in the genome-wide datasets. To do so, we trained linear classifiers using the most variable genes to distinguish perturbed cells from non-targeting controls, evaluating performance on held-out cells corresponding to the same perturbation (**Methods**). Our analysis indicates that a considerable proportion of on-target perturbations in expressed genes result in discernible single-cell transcriptomic changes (48.36% in HCT116, 40.37% in HEK293T, compared to 54.30% in K562 ^17^, **Figure 3A**). This suggests that numerous genetic perturbations induce detectable transcriptomic phenotypes. Classification results are not driven by expression of the targeted primary transcript, as target-ablated results look similar to non-ablated ones (**Figure S2A**). In contrast, classification of non-targeting perturbations as a negative control yielded markedly fewer discernible transcriptomic changes (4.8% in HCT116 and 0.0% in HEK293T cells; **Figure S2B-C**). This analytical approach may identify non-targeting sgRNAs with potential off-target effects – similar to those observed with CRISPRi perturbations (e.g. Figure S4G-H in Replogle et al. 2022 ^17^) – allowing their exclusion from analyses or future library designs.

**Figure 3.**
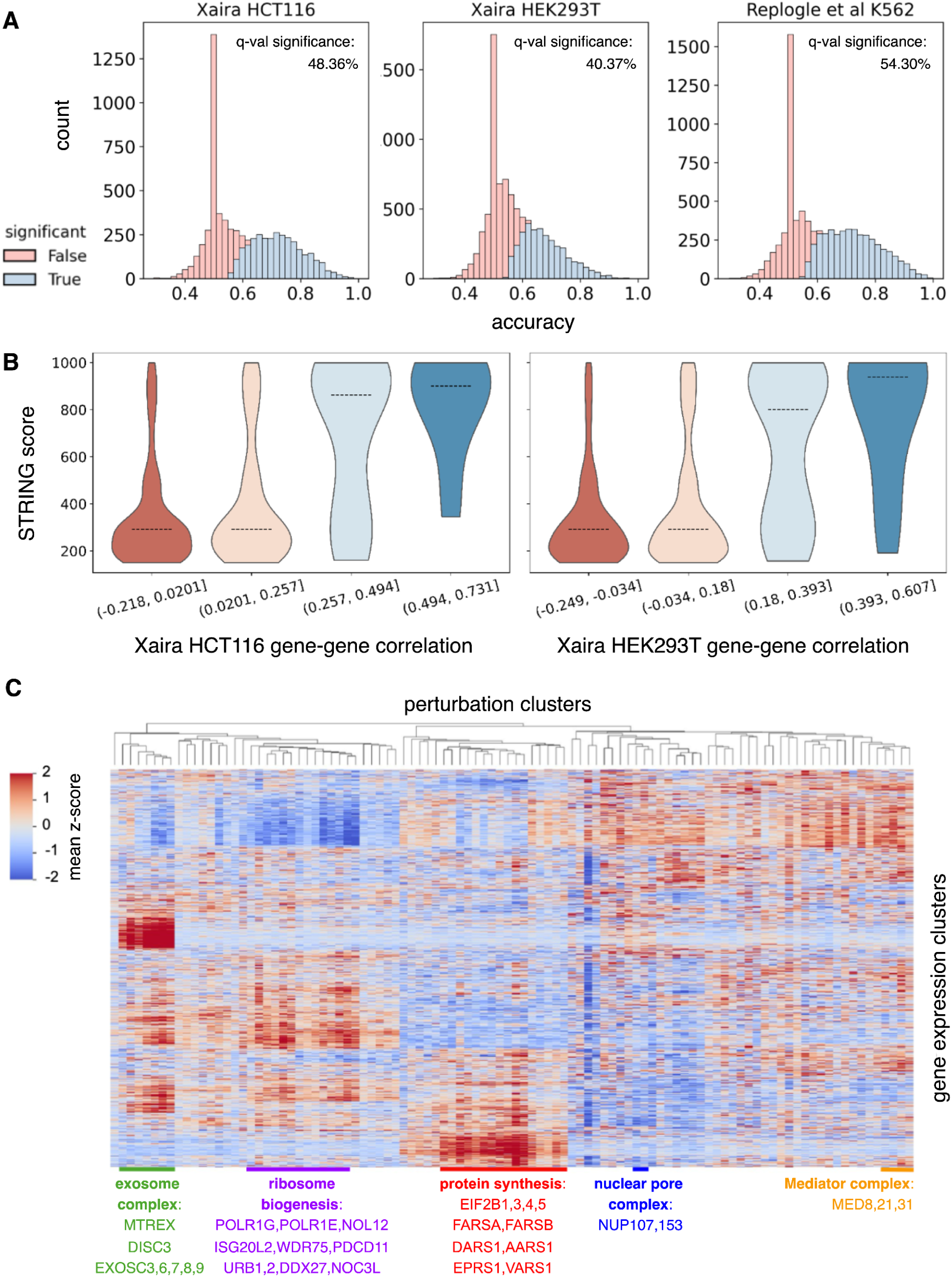
FiCS Perturb-seq captures biologically meaningful transcriptomic changes. **A**. Distributions of perturbation-level binary classifier accuracies across datasets. Significance was determined using the Storey q-value method ^41^. **B**. STRING scores of gene-gene pairs divided into bins based on their expression profile correlation. Dotted line indicates median STRING score in each bin. **C**. Heatmap of perturbations with strong transcriptional phenotypes clustered by expression of highly variable genes. Perturbation clusters are labeled with manual annotations.

To evaluate whether our transcriptomics-derived network aligns with known biology, we systematically compared it against physical StringDB — a curated database of known gene-gene interactions derived from biochemical studies of physical protein complexes ^32^. We computed gene-gene correlations within our perturbation atlas, grouped these relationships by correlation strengths, and compared their strength against StringDB interaction scores (**Methods**). This analysis revealed that StringDB scores significantly increased with transcriptomic correlation strength (**Figure 3B**), suggesting a strong concordance between networks derived from Perturb-seq correlations and known protein interaction networks.

Next, we clustered perturbations with strong phenotypes based on their transcriptional responses (**Methods**), which effectively grouped genes from related functional pathways or protein complexes. For example, genes associated with ribosome biogenesis formed a distinct cluster (**Figure 3C**). This cluster included components of RNA polymerase I for rRNA transcription (POLR1G, POLR1E); factors involved in pre-rRNA processing (NOL12, ISG20L2); proteins essential for nucleolar ribosome maturation (WDR75, PDCD11, URB1, URB2, DDX27, NOC3L) and cytoplasmic ribosomal maturation (RSL24D1); and ribosomal proteins (RPL37A, RPL4) ^33^.

Other notable clusters included genes encoding components of the exosome complex, which is responsible for RNA degradation ^34^; the Mediator complex ^35^, the nuclear pore complex ^36^, and the eIF2B complex ^37^ and aminoacyl-tRNA synthetases ^38^, which are both essential for protein synthesis (**Figure 3C**). Additionally, we observed that perturbations of mitochondrial proteins (TIMM44, PHB1, SAMM50, and PRELID1) clustered with those of the eIF2B complex members (**Figure 3C**). This finding is consistent with recent reports that mitochondrial injury induces phenotypes resembling the integrated stress response ^17,39,40^.

### sgRNA abundance stratifies KD efficiency and detects dose-dependent genetic effects

Perturbed cells exhibited notable differences in median KD efficiency, the degree to which expression of a target gene is inhibited by the CRISPRi system, between cell lines: 75.4% in HCT116 versus 51.5% in HEK293T (**Table 1**, **Figure 4A**). As both cell lines received identical dCas9 and sgRNA delivery vectors, we hypothesized this variability arises from differing expression levels of key CRISPRi components. Several potential factors that are directly measured from Perturb-seq results include: 1) sgRNAs; 2) the scaffolding protein TRIM28; and 3) the KRAB-interacting silencing factor HP1a ^27,42^. We found that baseline expression levels of sgRNAs and TRIM28 were significantly higher in HCT116 compared to HEK293T (fold change = 2.26 and 2.61, Mann-Whitney U pval = 1.61e-32, 3.48e-150, **Figure 4B**). This suggests that the abundance of sgRNAs and TRIM28 may be rate-limiting for recruiting repressive factors and could predict CRISPRi performance across cell systems.

**Figure 4.**
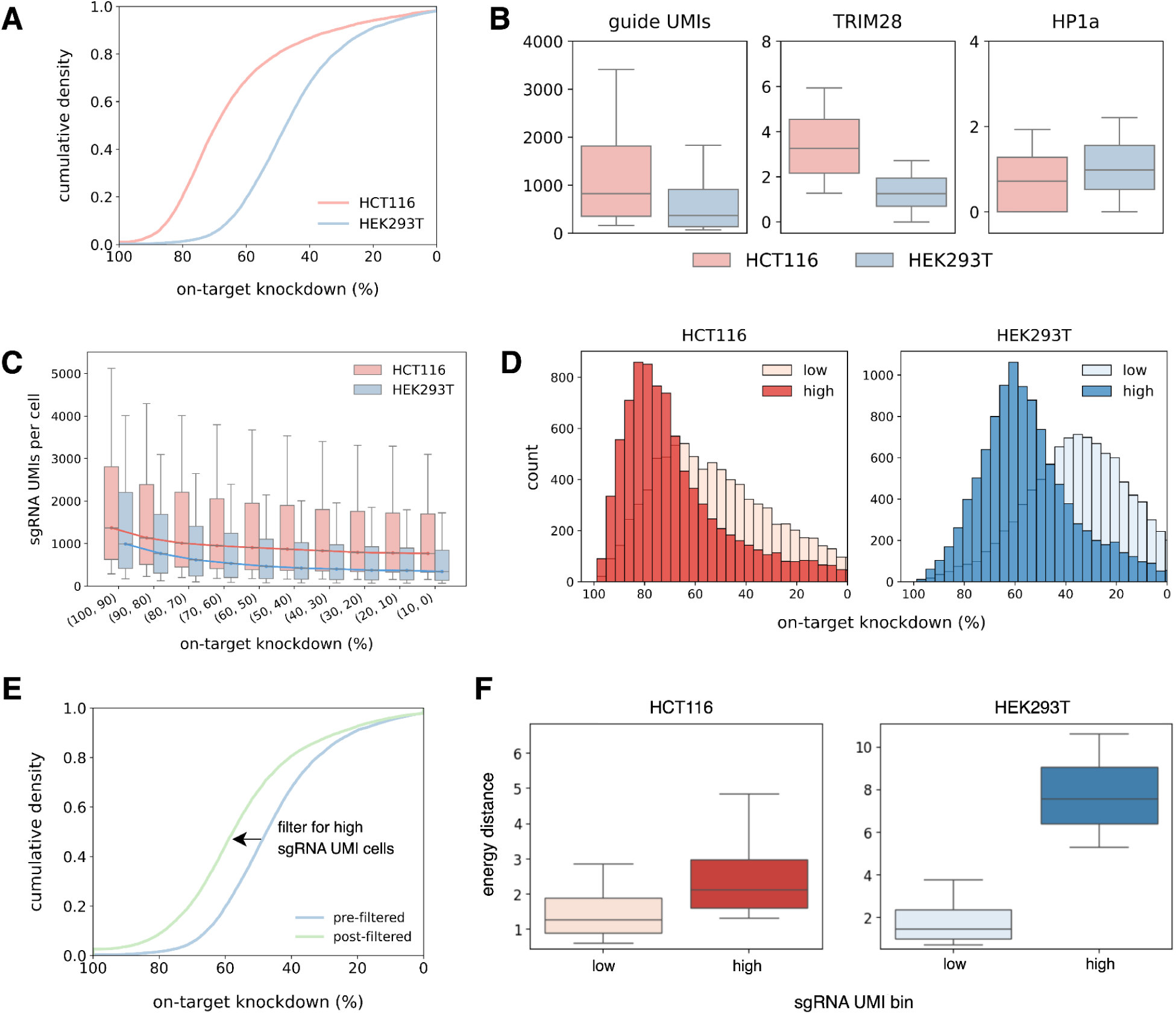
sgRNA UMI counts stratify KD efficiency and detect dose-responsive transcriptomic changes. **A**. Cumulative density function (CDF) of on-target KD percentage in expressed genes in HCT116 and HEK293T cells. **B**. Distribution of sgRNA UMIs, TRIM28, and HP1a counts in all HCT116 and HEK293T cells. Raw counts are shown for sgRNA UMIs. cp10k counts are shown for TRIM28 and HP1a. **C**. sgRNA UMI counts of cells binned by on-target KD of expressed genes in HCT116 and HEK293T. Line denotes median sgRNA UMI counts across KD bins. **D**. Distribution of on-target KD percentages in cells with low (bottom 25%) or high (top 25%) sgRNA UMI counts for cells containing perturbations targeting expressed genes. 15.8% and 4.2% of perturbations had 100% KD in low or high groups in HCT116 and HEK293T, respectively, likely due to sparsity of scRNAseq and were excluded from plotting. **E**. CDF of on-target KD efficiency of expressed genes in pre-filtered (not filtered by sgRNA UMI count) or post-filtered (filtered by sgRNA UMI count) HEK293T cells. **F**. E-distance in cells with low (bottom 25%) or high (top 25%) sgRNA UMI counts. **B,C,F**. Whiskers represent the 10th and 90th percentiles, while the box represents the 25th, 50th, and 75th percentiles.

In addition to inter-cell line variability, KD efficiency also varies considerably among cells receiving the same perturbation within a single cell line. This variability of KD could potentially be harnessed for studying dose dependent genetic effects and/or modeling heterogeneity within a cell population; however, KD levels in individual cells are difficult to quantify due to transcript dropout and data sparsity in scRNAseq data. While computational methods exist to infer perturbation strength from downstream transcriptional changes at the single-cell level (e.g., the perturbation-response score by Song et al., 2025 ^20^), we sought a more direct proxy related to the perturbation delivery itself. We noted that sgRNA transcripts, driven by a robust Pol III promoter, are abundantly detected by the “direct capture” approach (often hundreds of UMIs per cell). Therefore, we explored using sgRNA abundance as a proxy for inferring KD effectiveness. Indeed, we found a strong correlation between KD efficiency and sgRNA level per cell in HCT116 (R = 0.910, pval = 2.54e-4) and HEK293T (R = 0.901, pval=3.67e-4, **Figure 4C**). In addition, we showed that by binning cells into high (top 25%) and low (bottom 25%) categories based on their sgRNA UMI counts per perturbation, we could successfully stratify cells according to their KD levels (two-sample KS pval=9.08e-39 in HCT116 and 1.23e-100 in HEK293T, **Figure 4D**). Consequently, sgRNA counts proved effective as a filtering mechanism; selecting the top 25% of cells based on sgRNA expression enhanced the KD of HEK293T expressed genes (**Figure 4E**).

We next investigated whether stratifying cells based on this measure (sgRNA abundance) could not only effectively distinguish varying perturbation strengths but also delineate corresponding functional impacts. To this end, we focused on a subset of perturbations targeting essential genes, which typically induce strong transcriptomic changes ^17^. Indeed, the high-sgRNA group consistently yielded a higher energy distance (e-distance) between on-target perturbations and non-targeting control cells compared to the low-sgRNA group, in both HCT116 (fold change of median e-distance = 1.66, pval = 2.87e-31) and HEK293T (fold change of median e-distance = 5.21, pval = 2.86e-167, **Methods**, **Figure 4F**) ^17^. These results demonstrate that sgRNA abundance is a valuable instrument for stratifying KD efficiency, filtering out cells with weak perturbations, and enabling the detection of dose-dependent genetic effects.

## Conclusions

This work introduces FiCS Perturb-seq, an industrialized platform that successfully addresses critical bottlenecks in scalability and consistency that have previously constrained large-scale Perturb-seq studies. By strategically integrating reversible chemical fixation, fixation-compatible FACS enrichment, stable long-term cryopreservation, superloaded microfluidics, and extensive automation, FiCS Perturb-seq significantly enhances the practicality and reliability of generating genome-scale perturbation data. While our specific implementation utilized DSP chemistry and a reverse-transcription based workflow, the core principle of leveraging fixation suggests that similar advantages in mitigating batch effects and improving scalability could be realized with other fixation-based single-cell methodologies, such as FLEX (10x Genomics) ^43^.

The FiCS Perturb-seq platform is expressly designed to generate the scale and quality of data required to train biological foundation models. A key motivation for scaling – specifically, increasing cell coverage per perturbation – was the shift from viewing CRISPRi outcomes as binary (KD vs. non-KD) to recognizing a spectrum of KD efficiencies across cells. Effectively interrogating this spectrum requires a substantial number of cells per perturbation to achieve adequate statistical power and resolution. Crucially, we demonstrated that sgRNA abundance can serve as a reliable proxy for KD efficiency. This finding allows for the stratification of cellular responses, enabling researchers to either selectively focus on strongly perturbed cells, or to explore dose-dependent genetic effects with unprecedented clarity – a capability that complements computational methods aimed at inferring perturbation dosage from downstream signatures ^20^.

The ability to quantify and leverage this perturbation spectrum has profound implications for the development of more sophisticated foundation models. By incorporating perturbation strength as a continuous variable, these models may achieve a more nuanced understanding of causal genetic interactions, interpret complex biological phenomena like haploinsufficiency and compensatory mechanisms, and facilitate more quantitative, dose-aware approaches to target discovery and prioritization. The X-Atlas/Orion dataset provides a rich, publicly available resource to catalyze these advancements.

We envision this platform, and the foundational datasets it enables, will accelerate the construction of robust, causal foundation models, thereby significantly advancing the frontier of predictive biology.

## Methods

### Library design and cloning

In past genome scale Perturb-seq studies, an expression threshold cutoff has been used to identify genes to be included in the targeting library ^17^. In order to generate empirical data on the relationship between gene expression level and transcriptional effects, we included in our library design pairs of sgRNAs targeting every gene in the human genome ^44^.

KD perturbations were introduced using a dual sgRNA system driven by a single U6 promoter, separated by tRNA sequences that promote cleavage of the two sgRNAs ^28,29,45,46^. We modified pJR85 ^30^ to express mNeonGreen and puromycin resistance markers and drive expression of two S. pyogenes sgRNAs separated by one of four tRNA sequences. The top sgRNA pair for each gene was synthesized by concatenating: 5’ PCR handle sequence/BsmBI site – AGGCACTTGCTCGTACGACGCGTCTCACACCG, first protospacer, first sgRNA scaffold – GTTTCAGAGCTAAGCACAAGagtgcatagcaagttgaaataaggctagtccgtttacaacttggccgctttaaggccggtcctagcaaggccaagtggcacccgagtcgggtgc, tRNA separator, second protospacer, 3’ PCR handle sequence/BsmBI site – GTTTCGAGACGATGTGGGCCCGGCACCTTAA (Twist Biosciences).

The four tRNA sequences used were:

Alanine tRNA ^28^ AACAAAGGGGGTATAGCTCAGTGGTAGAGCGCGTGCTTAGCATGCACGAGGTCCTGGGTTCGATCCCCAGTACCTCCA

modified human cysteine tRNA ^47^ AGAGGGGGTATAGCTCAGTGGTAGAGCATTTGACTGCAGATCAAGAGGTCCCCGGTTCAAAT CCGGGTGCCCCCT

human glutamine tRNA ^46^ GGCGGTTCCATGGTGTAATGGTTAGCACTCTGGACTCTGAATCCAGCGATCCGAGTTCAAATCTCGGTGGAACCT

human glycine tRNA ^46^ ATCGGGCATGGGTGGTTCAGTGGTAGAATTCTCGCCTGCCACGCGGGAGGCCCGGGTTCGATTCCCGGCCCATGCA

Second-strand synthesis of oligonucleotide pool was performed with a primer

(5′-TTAAGGTGCCGGGCCCACATCGTCTCGAAAC-3′) at a primer-to-template molar ratio of 100:1. Primer annealing was carried out at 67.7 °C for 5 minutes, followed by a temperature hold at 4 °C. Subsequently, DNA Polymerase I, large (Klenow) fragment and dNTPs were added to the reaction mixture. Primer extension was conducted with an incubation at 25 °C for 15 minutes, followed by enzyme inactivation at 75 °C for 20 minutes, and a final hold at 4 °C. The resulting DNA was purified using SPRI bead-based cleanup. The purified insert pool was cloned into a linearized recipient vector using Golden Gate assembly. Post-assembly, plasmid DNA was purified via isopropanol precipitation and subsequently transformed into electrocompetent *E. coli*. The bacteria were propagated in liquid culture, and colony counts were performed to assess transformation efficiency. Final library plasmid pool purification was carried out using the Qiagen Plasmid Maxi Kit, in accordance with the manufacturer’s instructions (<u>Qiagen Manual</u>).

### Lentivirus Production

Post-cloning, the library plasmid pool was subjected to next-generation sequencing (NGS) library preparation to assess the overall distribution and skew of the library prior to transfection into lentivirus. Once the skew ratio (90th percentile counts / 10th percentile counts) was determined to be less than three, the pooled library was prepared for lentiviral transfection using the Lenti-X system, ViraPower™ Lentiviral Expression System, Lipofectamine 3000, and the purified plasmid. For large-scale lentivirus production, transfection was performed in 15 cm dishes, with Lenti-X cells plated at a density of 0.5 million cells/mL. Plasmids were thawed on ice prior to initiating the transfection process.

To prepare the DNA-Lipofectamine 3000 complexes for each transfection, Tube A was prepared by combining 125 µL Opti-MEM, 1.5 µg ViraPower Packaging Mix, 0.5 µg plasmid DNA, and 4 µL of P3000 reagent (added last) in the specified order. Tube B was prepared by combining 6 µL Lipofectamine 3000 and 125 µL Opti-MEM, vortexing the solution, and incubating for 5 minutes at room temperature (RT). After incubation, the contents of Tube A and Tube B were combined, mixed thoroughly, and incubated for an additional 20 minutes at RT to allow for complex formation. The scale of the reaction can be adjusted for larger production volumes.

For transfection, 250 µL of the DNA-Lipofectamine 3000 mixture was added dropwise to each well containing Lenti-X cells. The media was replaced with 2 mL of antibiotic-free DMEM supplemented with 10% fetal bovine serum 6–18 hours post-transfection.

Viral supernatants were collected 48–72 h post-transfection, with approximately 2 mL per well. The supernatant was filtered using a 0.45-µm PVDF filter and stored at −80°C, or alternatively, it was centrifuged at 3,000 rpm for 15 min at 4°C to pellet cellular debris before storage.

### Cell culture and cell line generation

HCT116 and HEK293T cells were propagated in RPMI 1640 GlutaMAX without HEPES (Gibco) supplemented with 10% fetal bovine serum and 100 units/mL of penicillin and 100 μg/mL of streptomycin. Both cell lines were stably integrated with a dCas9-mScarlet-ZIM3 based CRISPRi effector expressing a blasticidin resistance marker using PiggyBac transposon gene delivery according to the manufacturer’s instructions (SBI) and sorted to >95% purity for mScarlet fluorescence and cultured in medium containing 10 μg/mL for at least 5 days before transduction with the genome wide sgRNA library.

### Perturbation screening

For both screens, the following coverage calculations were used to determine cell number and viral amount: aiming for >1000-fold cell coverage at the viral transduction step at a multiplicity of infection of 0.1, 2.3e8 cells were transduced with 2.3e7 transducing units of genome-wide library lentivirus. Cells were transduced by adding processed viral supernatants to CRISPRi expressing cells and 8 µg/mL polybrene. Transduced cells were selected with 1 μg/mL puromycin days 3-5 post transduction and prepared for 10X single cell sequencing on day 7 post-transduction.

### Cell sorting, fixation and cryopreservation

For both screens, >2e8 cells were prepared for sorting by resuspension in ice cold DPBS at 1e6 cells/mL and stained with Zombie NIR dye at a 1:1000 dilution (BioLegend). Cells were fixed in a solution of 1 mg/mL of DSP (Thermo) in PBS for 30 min at room temperature in the dark. The fixation buffer was freshly prepared by diluting a 50 mg/mL stock DSP solution dissolved in DMSO dropwise into PBS and filtering through a 40 µM FlowMi strainer to remove DSP precipitate ^24,48^. After puromycin selection, cells were >70% double positive for mScarlet and mNeonGreen fluorescence. Viably fixed and perturbed cells were enriched by sorting on a SH800 (Sony) or CytoFlex (Becton Dickinson) sorter gated on Zombie NIR-negative, mScarlet-positive, mNeonGreen-positive single cells. Cells were sorted into pre-coated tubes containing a cell recovery buffer of 1% bovine serum albumin in ice cold PBS, supplemented with 1 U/μL of RNasin RNase inhibitor (Promega). Cells were pelleted and resuspended at 1e6 cells/μL in Bambanker cell freezing media according to the manufacturer’s instructions (Nippon Genetics Co.) supplemented with 0.2 U/μL RNasin and cryopreserved at −80°C until preparation for scRNAseq.

### Cell thawing and preparation for 10X scRNAseq

Prior to cell thawing, the necessary buffers were freshly prepared and maintained on ice. The 10x Cell Wash Buffer was prepared by combining 6.67 mL of 7.5% BSA solution with 43.33 mL of 1x DPBS, yielding a total volume of 50 mL. Additionally, the 10x Cell Loading Buffer with RNase Inhibitor was prepared by mixing 4,975 µL of 10x Cell Loading Buffer with 25 µL of RNase Inhibitor (40 U/µL), resulting in a final volume of 5,000 µL. These buffers were kept on ice throughout the procedure to maintain optimal conditions for cell processing.

Cells were rapidly thawed in a 37°C water bath until a small ice crystal remained (∼1 min). To prevent osmotic shock-induced lysis, 10 mL of ice-cold 10x Cell Wash Buffer (1x DPBS + 1% BSA) was added dropwise to the thawed cells in a 15 mL Falcon tube, followed by gentle mixing.

Cells were then centrifuged at 500 × g for 5 min at 4°C (acceleration = 9, deceleration = 9). The supernatant was carefully aspirated using a biosafety cabinet (BSC) vacuum line with a 2 mL serological pipette, ensuring minimal disturbance to the cell pellet. The final aspiration step was performed using a P1000 pipette to remove residual liquid.

A second wash was performed by resuspending the pellet in 1 mL of ice-cold 10x Cell Wash Buffer supplemented with 0.2 U/µL RNase inhibitor. The cell suspension was centrifuged again at 500 × g for 5 min at 4°C under identical acceleration and deceleration conditions. Following centrifugation, the supernatant was carefully removed using a P1000 pipette, leaving behind 200 µL of buffer. The pellet was gently resuspended to ensure a uniform single-cell suspension.

### Cell Counting & 10x GEM-X Single Cell 5’ v3

A 1:10 dilution was prepared by adding 1 µL of the cell suspension to 9 µL of 10x Cell Loading Buffer. From this dilution, 8 µL was mixed with 8 µL of trypan blue, and 10 µL of the mixture was counted on a hemocytometer. Following cell counting, the 10x Genomics Chromium GEM-X 5’ Chip was superloaded at 100,000 cells per lane to achieve a target recovery of approximately 70,000 cells per lane. The GEM generation, barcoding, and subsequent library preparation were performed according to the <u>10X Chromium GEM-X Single Cell 5’ v3</u> with Feature Barcode Technology for CRISPR Screening protocol. We generated the data for this manuscript using a combination of automated and manual library preparation methods.

### Automation

The following sections of the 10X GEM-X Single Cell 5’ v3 kit protocol were automated on the Hamilton Vantage liquid handler:

1. cDNA cleanup and size selection with SPRIselect
2. Gene Expression Library Construction

The following is the configuration of our dual arm Vantage liquid handler:

1. multiprobe 96 head (MPH96)
2. 4 x 1000uL single channels
3. 8 x Magpip channels
4. 4 x Alpaqua FLX plate magnets
5. 1 x Hamilton Heater Shaker (HHS)
6. 1 x Hamilton Heater Cooler (HHC)
7. 2 x Inheco On Deck Thermal Cyclers (ODTC) in the logistics cabinet
8. Custom cooler carrier that is connected to a chiller circulator that can cool up to 5 plates. We also built custom aluminum blocks to fit both deep well and PCR plates.
9. Track gripper to shuttle labware between the deck and the ODTC
10. Exit Entry module for expandability
11. HEPA hood and UV lamp

The automation method has a flexible throughput of one to four 96 well plates for full plate runs, or 12 to 84 samples for a partial plate operation.

We used the Hamilton Liquid Verification Kit (LVK) to define liquid classes that are involved in the 10X GEM-X workflow for both the Magpip channels and the MPH96 (**Table 2**). All liquid classes are developed with <3% CV and trueness (R%).

**Table 2.** 10x Genomics GEM-X Liquid Class Definition Performances. For the highest volume transfer reliability, we added specific correction curve points for each volume that is used in the 10x Genomics GEM-X workflow.

| Liquid | Pipette Type | Vol (uL) | CV% | R% |
| --- | --- | --- | --- | --- |
| AMP master mix | Magpip | 110 | 0.67 | 0.79 |
|  |  | 165 | 0.69 | 0.95 |
|  |  | 55 | 1.28 | 1.46 |
|  | MPH96 | 50 | 2.63 | 1.83 |
|  |  | 95 | 1.1 | 1.01 |
| DPBS | Magpip | 90 | 0.5 | 0.45 |
|  |  | 10 | 0.44 | 0.8 |
| DPBS | MPH96 | 10 | 0.37 | 0.5 |
|  |  | 90 | 0.09 | 0.42 |
| EB | MPH96 | 5 | 0.72 | 0.66 |
|  |  | 10 | 0.42 | 0.9 |
|  |  | 30 | 1.73 | 2.66 |
| Frag master mix | Magpip | 132 | 0.57 | 0.75 |
|  |  | 88 | 0.78 | 1.61 |
|  | MPH96 | 50 | 1.48 | 2.12 |
| Ligase master mix | Magpip | 55 | 0.88 | 0.21 |
|  |  | 110 | 1.63 | 0.51 |
|  | MPH96 | 50 | 1.22 | 1.11 |
| SPRI | MPH96 | 30 | 0.73 | 0.64 |
|  |  | 20 | 0.53 | 0.37 |
|  |  | 70 | 2.36 | 1.48 |
|  |  | 100 | 0.87 | 1.12 |
|  |  | 50 | 0.25 | 0.13 |
| SPRI supernatant | MPH96 | 75 | 1.09 | 0.36 |
|  |  | 150 | 2.34 | 0.87 |

### QC, Pooling & Sequencing

Sequencing was performed using an Illumina 25B NovaSeq X kit, with typically four to five Gene Expression libraries and four to five paired CRISPR libraries pooled per run. The sequencing strategy aimed to account for loading and counting errors, targeting a recovery of 70,000 cells per lane. The Gene Expression libraries were sequenced to a depth of 50,000 reads per cell, while the CRISPR libraries targeted 5,000 reads per cell per sample.

Final Gene Expression and CRISPR libraries were separately pooled to a 10 nM concentration, determined through high-sensitivity DNA Qubit quantification and TapeStation D1000 ScreenTape analysis, using the average fragment size. The TapeStation average fragment size was calculated using a range of 200 bp to 1000 bp. These 10 nM final libraries were then analyzed using the <u>CFX Opus 384 Real-Time PCR System (#12011452)</u> with the <u>Kapa Library Quantification Kit Universal qPCR Master Mix (KK4824)</u> to ensure precise quantification prior to sequencing.

After qPCR validation, the 10 nM Gene Expression and CRISPR pools were diluted with Qiagen EB buffer, adjusting to a final concentration of 2 nM with a 2% PhiX spike-in. The final 2 nM pooled libraries were then denatured and sequenced on the Illumina NovaSeq X (25B), following the <u>Illumina NovaSeq X protocol</u>, with a final loading concentration target of 170 pM. Sequencing parameters were set to 28/10/10/90, per 10x Genomics specifications.

### Basecalling, alignment, and sgRNA calling

bcl2fastq2 version 2.2 was used for basecalling and FASTQ demultiplexing. Cell Ranger version 8.0.1 was used for alignment, cell barcode calling, and UMI counting. The 10x Genomics GRCh38 2024-A pre-built genome was used as the reference transcriptome for alignment. After alignment, QC metrics, such as mean reads per cell, median genes per cell, and median CRISPR UMIs per cells, were examined for each GEM group to ensure sufficient sequencing depth and coverage.

Next, sgRNA calling was performed by aggregating the sgRNA UMI-cell barcode matrices across GEM groups. A two-component Gaussian mixture model was fit on each sgRNA’s data in the aggregated sgRNA UMI-cell barcode matrix to determine the minimum UMI threshold for each sgRNA across the entire dataset. The minimum UMI threshold was defined as the greater value between the user-specific minimum (in this case, 3) and the lowest UMI count assigned to the high-expression component of the Gaussian mixture model. Based on this threshold, each cell was assigned zero, one, or multiple sgRNA identities. Only cells with the expected two-sgRNA same-gene-target pair were used for calculating on-target KD in expressed genes and subsequent downstream analyses. Expressed genes were defined as genes that collectively account for 99% of the cp10k (counts per 10,000) counts averaged across cells containing a non-targeting sgRNA pair. 11,181 genes and 12,573 genes were defined as expressed in HCT116 cells and HEK293T cells, respectively.

### Batch-batch correlation of X-Atlas/Orion

To quantify batch-batch correlation, pseudobulked gene counts were obtained by summing raw counts of expressed genes from cells containing a non-targeting sgRNA pair. Pairwise Spearman correlations were then computed between selected sample groups within each dataset using pandas.DataFrame.corr(method=’spearman’). Sample groups were stratified based on median UMI counts, with the top and bottom quartiles randomly sampled to ensure balanced comparisons. Additionally, unpaired correlations were calculated by randomly pairing samples between HCT116 and K562 datasets such that the number of pairs were the same as comparisons within a dataset. The Wilcoxon rank-sum test was used to test whether the correlation distribution of one dataset was significantly greater than another using scipy.stats.ranksum(alternative=’greater’). The median absolute deviation was calculated by subtracting the median correlation value from the correlation distribution, and then taking the median of the absolute value of the resulting residuals.

### Binary classification

A binary classification pipeline was employed to systematically identify gene perturbations with detectable transcriptomic effects. For each perturbation with at least 100 unique cells, logistic regression with L1-penalization (LASSO) was fit to distinguish perturbed cells from a balanced cohort of non-targeting control (NTC) cells. Raw counts were cp10k normalized, log1p transformed, then scaled by their mean. Variance-based feature selection limited the set of predictors to the 5,000 most variable genes for each perturbation.

Models were trained on an 80-20% stratified train-test split. Regularization strength was optimized via cross-validation within the training set across a range of lambda values using LogisticRegressionCV from sklearn. Performance was evaluated on a held-out set of cells. Statistical significance (p-values) was assessed using a one-sided binomial test, comparing the observed accuracy against a null hypothesis of random performance (accuracy = 0.5).

To correct for multiple hypothesis testing, we utilized the Storey q-value method ^41^, which estimates the proportion of true null hypotheses by analyzing the distribution of observed p-values, and adjusts p-values accordingly to control the false discovery rate. Significance was defined as q-value < .05.

Binary classifiers, particularly logistic regression with L1-regularization, directly quantify the accuracy and statistical significance of transcriptional signatures of a perturbation, even when only a small subset of transcripts are affected. In contrast, e-distance methods, which average transcriptional effects of a perturbation across the entire genome, have limited sensitivity to detect sparse or subtle transcriptomic effects and require computing costly permutational resampling distributions to evaluate statistical significance.

### Correlation of gene-gene expression profiles and corresponding STRING scores

Perturbations with strong phenotypes were chosen based on the following criteria: 1) present in at least 25 cells; 2) at least 30% on-target KD; 3) significant q-value in binary classifier, and 4) non-zero counts in the training split of non-targeting control cells in the binary classifier. 1,574 and 1,944 perturbations met these filtering criteria in HCT116 and HEK293T, respectively, and were used for downstream analysis. For feature selection, perturbations were represented by their mean normalized expression profile across 2,000 highly variable genes. Highly variable genes were determined using scanpy.pp.highly_variable_genes(n_top_genes=2000, flavor=’seurat_v3’). Cells were aggregated into pseudobulk profiles for each perturbation using z-normalized counts, resulting in an m x n matrix, where m represents the number of features and n represents the number of perturbations. z-normalized counts were determined by calculating the mean (μ_NTC_) and standard deviation (σ_NTC_) of expression in cells containing non-targeting sgRNA pairs across all GEM groups and then computed using the following formula for each gene, z = (x - μ_NTC_) / σ_NTC_. Spearman correlations were then determined for all pairwise gene-gene combinations using pandas.DataFrame.corr(method=’spearman’). Physical protein links were downloaded from the STRING database (9606.protein.info.v12.0.txt.gz and 9606.protein.physical.links.v12.0.txt.gz), and mapped to the gene-gene correlation pairs. Gene-gene pairs were then split into 4 bins, such that each bin has the same width, and then the distribution of STRING scores for each bin was visualized using kernel density estimates with seaborn.violinplot(cut=0).

### Clustering of perturbations with strong phenotypes

To identify perturbations with strong transcriptional phenotypes, we measured the transcriptional effect of perturbations targeting expressed genes in HCT116 data. Data was preprocessed using the following steps: 1) raw counts were normalized to cp10k counts and log1p transformed; 2) dimensionality reduction was performed on NTC cells by standardizing the counts using sklearn.preprocessing.StandardScaler and reducing the dimensionality to 20 components using sklearn.decomposition.PCA(n_components=20). A random subset of 1,000 NTC cells was selected and transformed using the fitted PCA model. Perturbations targeting expressed genes and present in at least 100 cells were scaled and transformed into PCA space using the same scaler and PCA model fitted on the NTC data. The transcriptional effect for each perturbation was calculated using the formula: 2 * (mean Euclidean distance between perturbation and NTC) - (mean within-perturbation Euclidean distance) - (mean within-NTC Euclidean distance). The top 100 perturbations with the largest effect were selected for further analysis.

To identify highly variable genes across these perturbations, the transcriptomes of cells with the same perturbation were averaged to generate a mean transcriptome per perturbation using log2(cp10k) counts. The top 2,000 highly variable genes were identified from these averaged transcriptomes using sc.pp.highly_variable_genes(flavor=”seurat”). Z-scores were calculated across perturbations for each highly variable gene, resulting in a 100×2000 matrix. Z-scores were clipped to a range of −2 to 2 for visualization. Clustering and visualization of perturbations were performed using seaborn.clustermap(metric=“correlation”).

### Stratification by sgRNA and energy distance analysis

Following initial quality filtering of cells based on total counts, mitochondrial gene UMI percentage, gene counts, and guide detection, per-cell UMI counts for guide A, guide B, and combined guides were robust-z-scored within each GEM group to normalize sequencing depth variability. For each gene target separately, standardized total-guide UMI distributions were analyzed to categorize cells into “low” and “high” UMI groups. Specifically, cells with total-guide UMI z-scores in the lowest quartile were labeled as “low,” while those in the highest quartile were designated as “high,” effectively separating cells exhibiting particularly low or elevated UMI counts for subsequent downstream analysis.

Energy distance (e-distance) was calculated using the same method described in Replogle et al. 2022 ^17^. First, expression data was cp10k normalized, then scaled with z-scores, and transformed via PCA, fitting the PCA model to a combined reference group consisting of randomly sampled NTC cells plus high and low essential gene perturbation groups. Using coordinates from the top 20 PCs, e-distance scores were calculated for each perturbation group by comparing their PCA-derived distributions against the PCA coordinates of the NTC reference group. The e-distance metric quantifies differences between each perturbation group and the reference, capturing overall shifts or dispersion without assuming underlying distributions, thus capturing gross biological or technical differences between the groups.

## Supplemental Figures

**Supplemental Figure 1, related to Figures 1 and 2:**
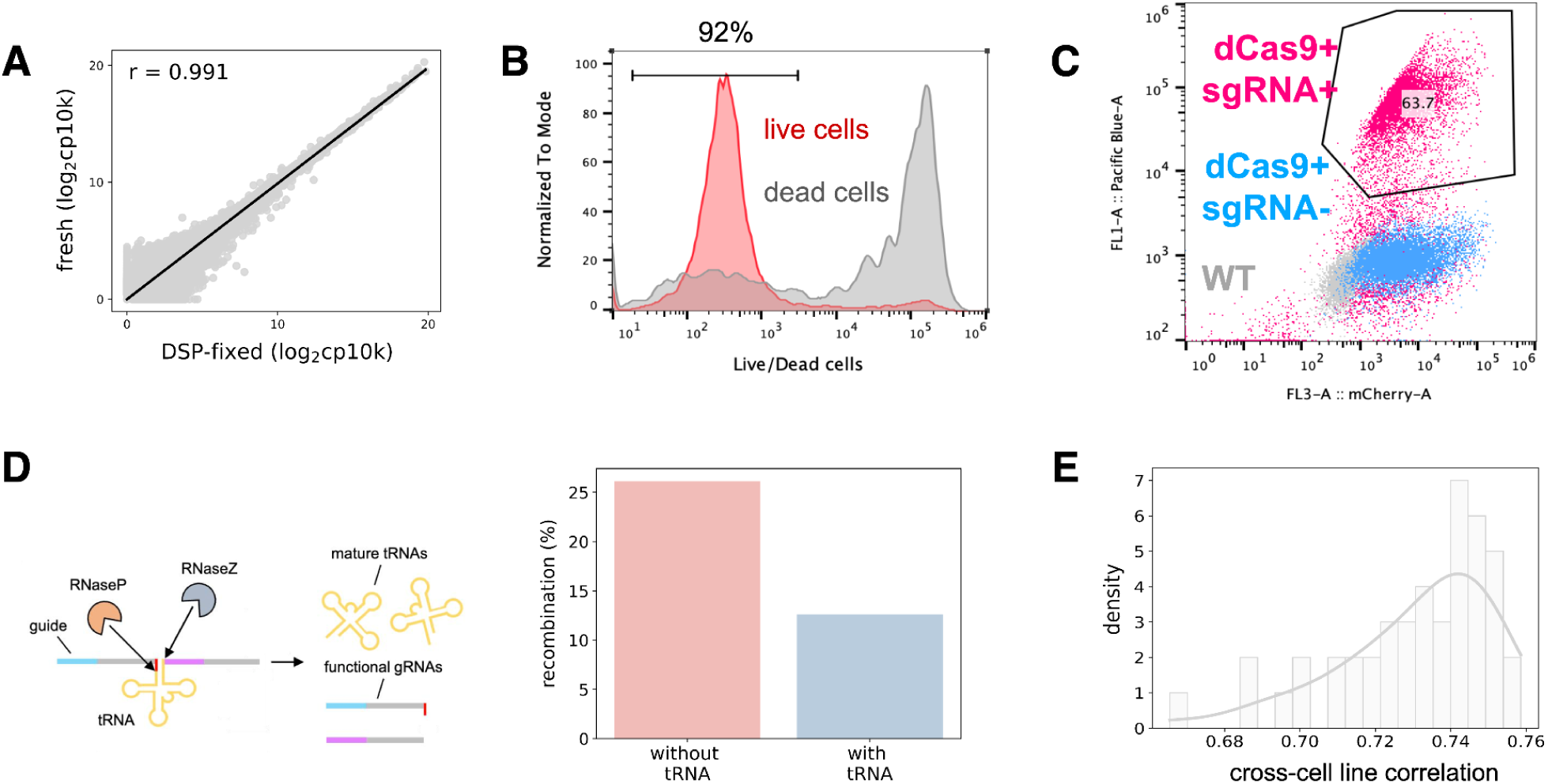
**A**: Scatterplot showing the correlation between pseudobulk counts from fresh and DSP-fixed HCT116 cells. Pseudobulk counts are calculated as log2(cp10k + 1), where cp10k represents counts per 10,000, with a pseudocount of 1. **B, C**: Flow cytometry plots from HCT116 showing the separation of live and dead cells using Zombie dye (**B**) and the enrichment of cells with high expression of dCas9 and sgRNA by double-sorting for mCherry/mScarlet (dCas9) and BFP/mNeonGreen (sgRNA) (**C**). **B**: 92% of live cells are negative for Zombie dye and sorted for downstream processing. **C**: 63.7% of live cells are double-positive for dCas9 and sgRNA and cryopreserved until library preparation. **D**: Schematic illustrating tRNA-based lentiviral delivery (**left**) and the percent recombination of cells without tRNA (26.1%) or with tRNA (12.6%) in HCT116 (**right**). Percent recombination was calculated as (number of two-guide cells - number of two-guide cells with the expected guide pair) / (number of two-guide cells) * 100. **E**: Intra-batch Spearman correlation of pseudobulked gene expression counts from cells with non-targeting sgRNAs in randomly paired batches from Xaira HCT116 and Replogle K562.

**Supplemental Figure 2, related to Figures 3 and 4:**
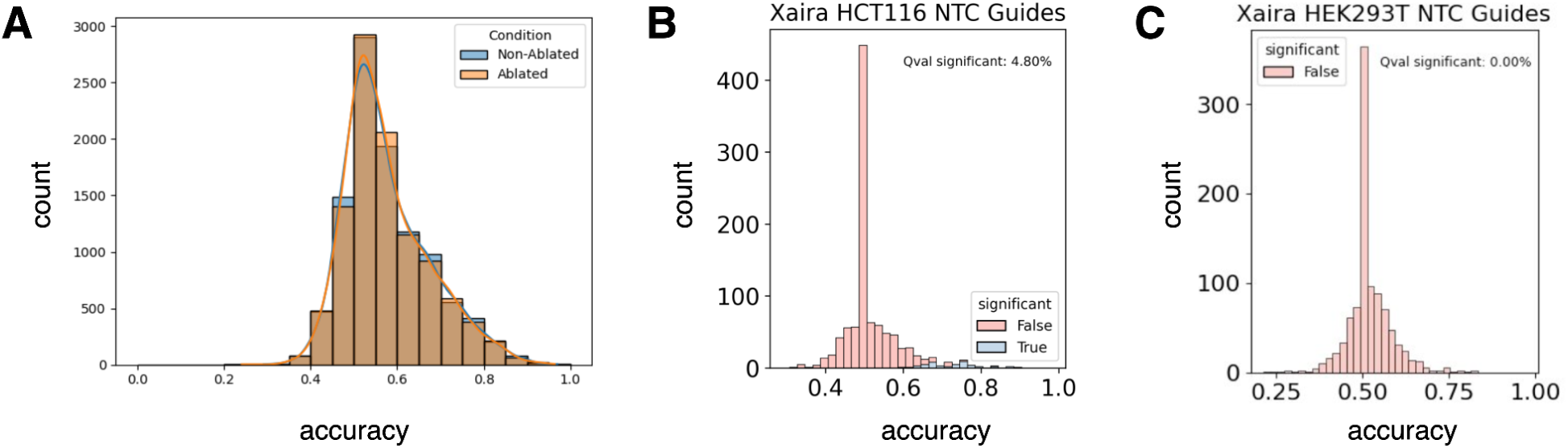
**A**: Distribution of classifier accuracy scores from perturbations when target is not ablated (blue) or ablated (orange) from transcriptome in HCT116. **B-C**: Distribution of classifier accuracy scores from perturbations targeting non-targeting perturbations in HCT116 (**B**) and HEK293T (**C**). Significance was determined using the Storey q-value method ^41^.

## Acknowledgements

We thank Marc Tessier-Lavigne for his valuable contributions to the dose dependency analysis and insights into viability sorting post-fixation. We also thank Jen Ong, Fudong Yin, Elise Ruark, and Brian A. Fox for their assistance with Figshare contracting and data upload. Lastly, we are grateful to Jonathan S. Weissman, Reuben A. Saunders, William E. Allen, Brian A. Fox, Nader Alerasool, Bo Wang, and Debbie Law for their helpful discussions and manuscript edits.

## Data Availability

h5ad files for cells with two sgRNAs targeting the same gene and from the same guide pair are available through FigShare under the following DOI: https://doi.org/10.25452/figshare.plus.29190726. h5ad files for all aligned cells will be released on Figshare at a later date.

